# Soybean CHX protein GmSALT3 confers leaf Na^+^ exclusion via a root derived mechanism, and Cl^−^ exclusion via a shoot derived process

**DOI:** 10.1101/2020.01.06.896456

**Authors:** Yue Qu, Rongxia Guan, Jayakumar Bose, Sam W. Henderson, Stefanie Wege, Lijuan Qiu, Matthew Gilliham

**Author notes:** These authors have contributed equally to this work (Yue Qu, Rongxia Guan). Funding information Natural Science Foundation of China, grant number 31830066; Scientific Innovation Project of Chinese Academy of Agricultural Sciences; ARC Centre of Excellence funding CE140100008; ARC Fellowships FT130100709, DE160100804 and DE170100346.

## Abstract

Soybean (*Glycine max*) yields are threatened by multiple stresses including soil salinity. *GmSALT3* confers net shoot exclusion for both Na+ and Cl^−^ and improves salt tolerance of soybean; however, how the ER-localised GmSALT3 achieves this is unknown. Here, GmSALT3’s function was investigated in heterologous systems and near-isogenic lines that contained the full-length *GmSALT3* (NIL-T; salt-tolerant) or a truncated transcript *Gmsalt3* (NIL-S; salt-sensitive). GmSALT3 restored growth of K+-uptake-defective *E. coli* and contributed toward net influx and accumulation of Na+, K+, and Cl^−^ in *Xenopus laevis* oocytes, while *Gmsalt3* was non-functional. A time-course analysis of the NILs confirmed that shoot Cl^−^ exclusion breaks down prior to Na+ exclusion, while grafting showed that shoot Na^+^ exclusion occurs via a root xylem-based mechanism. In contrast, NIL-T plants exhibited significantly greater Cl^−^ content in both the stem xylem and phloem sap compared to NIL-S, indicating that shoot Cl^−^ exclusion likely depends upon novel phloem-based Cl^−^ recirculation. NIL-T shoots grafted on NIL-S roots contained low shoot Cl^−^, which confirmed that Cl^−^ recirculation is dependent on the presence of GmSALT3 in shoots. Overall, these findings provide new insights on GmSALT3’s impact on salinity tolerance and reveal a novel mechanism for shoot Cl– exclusion in plants.

**Highlight:** GmSALT3 improves soybean salt tolerance. Here, using heterologous expression, we found GmSALT3 is a functional ion transporter, and, *in planta* that it confers shoot salt exclusion through root-based Na^+^ xylem exclusion and shoot-based Cl^−^ exclusion via phloem derived Cl^-^ recirculation.

## Introduction

There is a large market demand for soybean (*Glycine max*) as a staple edible oil and food crop; soybean is also an important component of intercropping and sequential cropping systems (Singh, 2010). As such, soybean production demand has been projected to increase by nearly 80 percent to 390 Mt by 2050 (Alexandratos & Bruinsma, 2012). However, salinity stress threatens soybean production. A soil salinity of EC (electrical conductivity) 18–20 dS/m, which is approximately 180 – 200 mM NaCl, has been shown to reduce yield by 61% from 2261 ± 438 kg/ha under control conditions to 881 ± 260 kg/ha (Chang, Chen, Shao, & Wan, 1994). In another study, Blanco et al. (2007) found that the emergence of soybean (cv. Conquista) was completely inhibited when EC was close to 8.0 dS/m. Yet, natural variation in soybean salinity tolerance has been observed, with some rare soybean germplasm found to be relatively salt-tolerant. In a large-scale evaluation of soybean salinity tolerance, less than 3% of the 10,128 evaluated soybean germplasm exhibited salt tolerance at both the germination and seedling stage; only 83 soybean cultivars were found to have significant tolerance to salinity at the vegetative stage (Shao, Wan, Chang, & Chen, 1993). It has been observed that salt-tolerant soybean cultivars often have superior agronomic performance to salt-sensitive cultivars (Phang, Shao, & Lam, 2008), indicating breeding for stress tolerance in soybean may improve overall yields even under standard conditions.

In soybean, a major quantitative trait locus (QTL) for salinity tolerance (*Ncl*) has been consistently mapped to chromosome 3 (Abel, 1969; Ha et al., 2013; Hamwieh et al., 2011; Hamwieh & Xu, 2008; Lee et al., 2004). Since 2014, several studies have been published focusing on the same dominant gene within this QTL on chromosome 3, in both wild (*GmCHX1*) and cultivated soybean (*GmNcl*/*GmSALT3*) (Do et al., 2016; Guan et al., 2014; Qi et al., 2014). This gene, denoted as *Glyma03g32900* in the reference soybean genome of the William 82 variety (William 82. a1.v1.1), is the premier candidate for conferring improved salt tolerance in the *Ncl* locus. It should be noted that in the William 82 genome a2.v1, *Glyma03g32900* appears to be incorrectly annotated as two transcripts (Glyma.03g171600 and Glyma.03g171700) rather than the full-length coding sequence obtained consistently from cDNA and encoding a full-length CHX protein. Here, we refer to the full-length gene obtained as *GmSALT3*.

According to phylogenetic analyses, the full-length GmSALT3 protein is a member of the CPA2 (Cation/Proton Antiporter 2) family of transporters; the most closely related sequence in *Arabidopsis thaliana* being AtCHX20 (a Cation/Proton Exchanger) (with 59% identity and 72% similarity) (Guan et al., 2014; Padmanaban et al., 2007). Several Arabidopsis CHXs, including AtCHX20, have been functionally characterised in heterologous systems; however, a connection to their function to roles in plants has been challenging, as examples of phenotypes associated with loss of function mutations is sparse (Sze & Chanroj, 2018). Functional studies of AtCHX have indicated that they might play role in modulating cation and pH homeostasis within the endomembrane system (Chanroj et al., 2011; Czerny et al., 2016; Padmanaban et al., 2007). The endomembrane localised AtCHX20 was, for example, suggested to be a K^+^ transporter involved in the osmoregulation of guard cells (Padmanaban et al., 2007).

In soybean, the salt-tolerant soybean cultivar Tiefeng 8 contains a full-length *GmSALT3*, while the salt sensitive cultivar 85-140 contains a 3.78-kb copia retrotransposon insertion in exon 3 of *GmSALT3* that truncates the transcript with a premature stop codon (making the 85-140 essentially a *Gmsalt3* knockout). This retrotransposon insertion resulted in the loss of the 10^th^ predicted transmembrane domain and the C-terminus of *GmSALT3* (Guan et al., 2014). *GmSALT3* is predominantly expressed in cells associated with the phloem and xylem (Guan et al., 2014); and expression of the full-length *GmSALT3*, but not the truncated version, results in low shoot Na^+^ and Cl^−^ accumulation in saline conditions (Guan et al., 2014; Liu et al., 2016; Qi et al., 2014). This low shoot NaCl accumulation is associated with increased crop yield (Do et al., 2016; Liu et al., 2016). The GmSALT3 protein has been localized to the endoplasmic reticulum (ER), sharing the membrane localisation of AtCHX20 in guard cells. This differs from most other transport proteins known to be involved in shoot Na^+^ and Cl^−^ exclusion, which are mainly localised on the plasma membrane or tonoplast (Munns and Gilliham, 2015).

Shoot salt exclusion requires the co-ordinated activity of many ion transporters that may: 1) limit the net entry of salt into the roots, 2) compartmentalise salt in the roots; and or 3) regulate vascular transport (most notably through root-based xylem exclusion) (Munns et al. 2020). Examples of transporters that have been implicated in salt tolerance in soybean include: the tonoplast localised GmNHX1 (Na^+^/H^+^ antiporter 1) (Li et al., 2006a), the plasma membrane localised GmSOS1 (salt overly sensitive 1, Na^+^/H^+^ antiporter) and the plasma membrane localised GmCAX1 (Ca^2+^/H^+^ antiporter 1) (Luo et al., 2005; Phang et al., 2008). These transporters are proposed to be involved in net exclusion of salt entry into roots (GmSOS1), salt compartmentation (*GmNHX1*) or signalling (*GmCAX1*). GmSALT3, was the first example of a transporter contributing to reduced net transfer of salt to the shoot in soybean (Cao, Li, Liu, Kong, & Tran, 2018). In other crop species, modulation of vascular ion load is commonly found to be one of the major salt tolerance mechanisms. For instance, in wheat, rice and Arabidopsis the HKT1;5-like (high affinity K^+^ transport) proteins are expressed in xylem parenchyma cells and facilitate Na^+^ retrieval from xylem sap back into the root (Munns et al., 2012; Møller et al., 2009; Ren et al., 2005; Uozumi et al., 2000). GmSALT3 differs from these previous proteins; the ER-localised is unlikely to directly retrieve from or load ions into the xylem sap. Therefore, GmSALT3 confers a novel salt tolerance mechanism which is yet to be determined.

Here, we investigated the function of GmSALT3 in heterologous systems and *in planta* to gain insights into the mechanisms by which GmSALT3 contributes to salt tolerance. In heterologous systems, *GmSALT3* complemented K^+^ uptake deficient *Escherichia coli*; while in *Xenopus laevis* oocytes, *GmSALT3* expression resulted in increased net influx of Na^+^, K^+^ and Cl^−^. Phenotypes of soybean NILs differing in *GmSALT3* alleles revealed that plants containing full-length *GmSALT3*, compared to truncated *Gmsalt3*, employ two distinct mechanisms to achieve shoot Na^+^ and Cl^−^ exclusion by facilitating net xylem retrieval of Na^+^ and a novel mechanism of greater phloem Cl^−^ retranslocation from shoots to roots. As such, we propose that the endomembrane GmSALT3 has a complex physiological role which ultimately impacts multiple ion fluxes across the plasma membrane.

## Materials and Methods

### Plant materials and growth conditions

Soybean plants were grown in a greenhouse (28 °C day, 16 h light and 25 °C night) at the Plant Research Centre, Waite Campus, University of Adelaide, Australia. Soybean seeds (Tiefeng 8, salt-tolerant parent; 85-140, salt-sensitive parent; NIL-T, near isogenic line with *GmSALT3*; NIL-S, near isogenic line with *Gmsalt3*) were imported from China through quarantine. Soybean seeds were germinated in pots containing a 1:1 mixture of perlite:vermiculite as described by Obermeyer & Tyerman (2005), with no nutrient solution added.

### cDNA cloning and plasmid preparation

To synthesize *GmSALT3* and *Gmsalt3* cDNA (2436 and 1131 nucleotides, respectively), total RNA was isolated from roots of 4-week old soybean plants using the TRIzol method (Shi & Bressan, 2006). First-strand cDNA was synthesized using a Thermoscript RT III kit (Invitrogen, USA). Gene specific primers (GmSALT3_gF and GmSALT3_gR, Gmsalt3_gF and Gmsalt3_gR; Table S1) were used to amplify *GmSALT3* cDNA with Phusion® High-Fidelity DNA Polymerase (New England Biolabs) using 35 cycles (98 °C 30 s, 65 °C 30 s, and 72 °C 150 s). Gel-purified PCR products were A-tailed using Taq polymerase (New England Biolabs) for 30 min at 72 °C, and then recombined into Gateway® entry vector PCR8/GW/TOPO (Invitrogen) for subsequent applications. Resulting clones were sequenced using internal primers (GmSALT3_F1, R1, F2, F3, F4, and F5, Gmsalt_F1, F2, and R1; Table S1). The sequence of *GmSALT3-YFP* was PCR amplified from a previously constructed plasmid *pBS-35S::GmSALT3-YFP*; the sequence of *GmSALT3-TM10* (first 10 transmembrane domain of *GmSALT3*) was PCR amplified from *PCR8-GmSALT3*. Gel-purified PCR products of *GmSALT3-YFP, GmSALT3-TM10* were recombined into entry vector PCR8 as described above.

*GmSALT3* and *Gmsalt3* CDS, *GmSALT3-YFP, GmSALT3-TM10, AtCHX20 and AtKAT1* within entry vector PCR8 were cloned into the pGEM-HE vector and a modified version of pET-DEST42 vector using Gateway ^®^ LR Clonase^®^ (Invitrogen, USA), for *X. laevis* oocyte expression and *E. coli* expression, respectively. The *E. coli* strain TK2463 lacks the T7 RNA polymerase, therefore, the T7 promoter in pET-DEST42 was replaced with the TAC promoter. Inverse PCR was performed with 5 ng of plasmid, 50 nM of forward and reverse primer (SH_215 and SH_216, Table S1), and 0.25 units Phusion High Fidelity Polymerase (New England Biolabs) in a final volume of 25 µL. Reactions were treated with Cloning Enhancer (Clontech Laboratories Inc) to remove circular template, and the plasmid was re-circularised using In-Fusion HD enzyme (Clontech Laboratories Inc).

### Growth assay in Escherichia coli

Competent cell preparation and *E. coli* transformation were conducted as described by Chanroj et al. (2011). The resulting transformants were grown on KML media (10 g/L tryptone, 5 g/L yeast extract, 10 g/L KCl) supplemented with 100 μg/mL carbenicillin and incubated at 37 °C overnight; positively transformed colonies were identified by colony PCR with gene specific primers (GmSALT3_gF and _gR; Gmsalt3_gF and gR). For *E. coli* growth experiments, freshly transformed cells were first grown overnight in 5 ml KML at pH 7.2. Cell cultures were replenished (OD_600_ = 0.5) and grown for 3 h in KML, and then washed with low potassium media (10 g/L tryptone, 2 g/L yeast extract, 100 mM D-mannitol) three times. Cells were normalized to OD_600_ 0.5 for a 96-well plate assay. In each well, 20 μL of normalized cell suspension was added to 180 μL growth solutions. All test media were supplemented with 100 μg/mL carbenicillin, with or without 0.5 mM IPTG (Isopropyl β-D-1-thiogalactopyranoside). The 96-well plates were inserted into FLUOstar Omega Fluorescence microplate reader (BMG LABTECH) set to 37 °C and A_600_ was measured every 15 min for 18 h with continuous shaking at 200 rpm.

### Characterization of GmSALT3 in X. laevis oocytes

pGEMHE-DEST containing *GmSALT3* was linearized using SphI-HF (New England BioLabs); cRNA was synthesized using mMESSAGE mMACHINE T7 Kit (Ambion). 46 nL/23 ng of cRNA or equal volumes of RNase-free water were injected into oocytes with a Nanoinject II microinjector (Drummond Scientific). Oocytes were incubated for 48 h in Calcium Ringer’s solution (96 mM NaCl, 2 mM KCl, 5 mM MgCl_2_, 5 mM HEPES, 0.6 mM CaCl_2_). All solution osmolalities were measured with a Vapor pressure osmometer (Wescor) and adjusted to 240–260 mOsmol kg^−1^ using mannitol. Measurement of ion profiles in oocytes followed Munns et al. (2012) with the following modifications. Six replications of 3 grouped oocytes were used for flame photometry (Sherwood 420), and Cl^−^ was measured by manual titrimetric Cl^−^ measurement using a chloride analyser (Sherwood® Chloridometer 926S). For YFP imaging, oocytes were analysed with a Zeiss LSM 5 Pascal Confocal Laser-Scanning Microscope equipped with an argon laser. Excitation/emission conditions for YFP were 514 nm/523-600 nm.

### Ion-selective microelectrode flux measurements from oocytes

The MIFE™ (Microelectrode Ion Flux Estimation technique; University of Tasmania, Hobart, Australia) allows noninvasive concurrent quantification of net fluxes of several ions using protocols outlined by Shabala et al. (2013) with oocytes being used rather than plant tissues in our experiments. Potassium and chloride commercial ionophore cocktails (99311, and 99408, respectively; all from Sigma-Aldrich) were used as LIX (Liquid Ion Exchanger) when measuring K^+^ and Cl^−^ fluxes. The Na^+^ fluxes were measured using an improved Na^+^ LIX described in Jayakannan et al. (2011). Oocytes were washed in ND96 (96 mM NaCl, 1 mM KCl, 1 mM MgCl_2_, 5 mM HEPES, pH 7.5). After incubation in ND96 for 72 hours, fluxes were measured after being transferred to BSM (5 mM NaCl, 0.2 mM KCl, 0.2 mM CaCl_2_, 5 mM HEPES, pH 7.5). During measurements, the ion-selective electrodes were positioned using a 3D-micromanipulator (MMT-5, Narishige, Tokyo, Japan), 100 μm from the oocyte surface. A computer-controlled stepper motor moved the electrode between two positions (100 and 200 μm, respectively) from the oocyte surface in 6 s cycles. The CHART software (Newman, 2001) recorded the potential difference between the two positions and converted them into electrochemical potential differences using the calibrated Nernst slope of the electrode. Net ion fluxes were calculated using the MIFEFLUX software for spherical geometry (Newman, 2001).

### TEM (Transmission Electron Microscopy)

Fresh soybean NIL-T and NIL-S roots were sectioned to 1 mm (length), stems were cross sectioned (1 mm in length and 1 mm in radius). Root and stem samples were incubated overnight in 1.5 ml microcentrifuge tubes with fixative (2.5% Glutaraldehyde, 4% Formaldehyde, 4% Sucrose, 0.1 M Phosphate buffer). Samples were washed three times in 1X PBS and then washed with osmium (263257, Sigma-Aldrich) for 4 hours. Sections were then washed three times in 1X PBS and soaked in 1X PBS for 20 mins and then embedded in a 1% agarose gel. Agarose embedded soybean root were cut into blocks (1 cm long). A series of dehydration steps were performed after embedding, using increasing concentrations of ethanol: 50%, 70%, 90% and 95% (each for 30 mins), and 100% ethanol for overnight dehydration. Samples were then incubated in increasing concentrations of resin (Spurrs): 5%, 10%, 15%, 20%, 25%, 30%, 40%, and 50% (each for 1 h), 50% overnight, 75% (4 h), 100% (4 h), and 100% (overnight). Resin wells containing the agarose blocks were then incubated at 60 °C for 3 days for polymerization. Polymerised resin capsules were cut to 70 nm thickness using a diamond knife and visualized under transmission electron microscopy (TEM; PHILIPS CM100).

### NaCl treatment and ion accumulation

Soybean plants were treated with 100 mM NaCl every 2 days (1.5 L solution per tray) at 14 days after sowing; the second dose of 1.5 L 100 mM NaCl was applied 16 days after sowing; the same volume of RO (reverse osmosis) water was applied to the tray of the control plants. Plant tissues were harvested and dried in an oven overnight at 60°C or freeze-dried overnight. Dry weight was recorded, then dried samples were digested in 10 mL 1% (v/v) nitric acid overnight at 65°C or freeze-thawed in Milli-Q water 3 times. Oven-dried samples were utilized to test Na^+^ and K^+^ accumulation with flame photometry (Sherwood 420), Cl^−^ was measured by titrimetric Cl^−^ measurement by using a chloride analyser (Sherwood^®^ Chloridometer 926S).

Freeze-dried samples were used to measure NO_3_ accumulation. The measurement method was modified according to Qiu, Henderson, Tester, Roy, & Gilliham (2016). Supernatant (50 μL) of freeze-dried samples in water was added into 200 μL of 5% (W/V) salicylic acid/H_2_SO_4_ and incubated at room temperature for 20 min, then 50 μL of this mixture was transferred into 950 μL of 2 M NaOH and incubated at room temperature for at least 20 min (to cool down to room temperature). Of this, 250 μL was loaded into separate wells of a flat-bottom transparent 96-well plate (Greiner Bio-One, Austria). Absorbance was measured at 410 nm. Standards measured included 0, 0.5, 1, 2, 2.5, 5, 7.5, and 10 mM KNO_3_.

### Phloem and xylem sap extraction

Phloem sap was extracted according to the method of Rupassara (2008) and Ren et al. (2005). Soybean and stems were cut approximately 2 cm above the ground level with the upper part immediately dipped in 1.5 mL of 0.1 mM EDTA solution (pH adjusted to 8 with NaOH) in a 2 mL microcentrifuge tube for 20 min, then the tube was immediately dipped in liquid nitrogen and stored in -80°C until analysis. The glutamine concentration in the extract was measured using a Glutamine Assay Kit (EGLN-100, EnzyChrom™, BioAssay Systems). Xylem sap was extracted using a pressure chamber. Lower parts of the cut stems with roots were transferred into the pressure chamber. Pressure was increased gradually (0.05 MPa increments) until the xylem sap presented. The first two drops emerging were discarded using a micropipette to reduce contamination from damaged cells or phloem sap (Berthomieu et al., 2003). Xylem sap was then collected during the following 5 min, and stored at -20 °C until analysis. Ion contents was measured as stated above.

### Grafting of soybeans

NIL-T and NIL-S were grown in a growth chamber with a 16 h light (28 °C)/8 h dark (25 °C) cycle at 60% humidity. The grafting protocol followed Guan *et al*. (2014) with the following modifications. After grafting, when the first trifoliate leaf fully expanded (about 10 days after grafting), healthy plants were selected and treated with 100 mM NaCl for 8 days. Na^+^, K^+^, and Cl^−^ contents were measured in soybean leaves and roots.

## Results

### GmSALT3 expression impacts Na^+^, K^+^, and Cl^−^ fluxes in heterologous systems

We functionally characterised GmSALT3 in heterologous systems, including *E. coli* and *X. laevis* oocytes. The *E. coli* strain TK2463 (*trkAΔ, kup1Δ, kdpABCDEΔ*) is defective in its native K^+^ uptake mechanisms (Epstein, Buurman, McLaggan, & Naprstek, 1993) (Fig. 1) and shows a reduced growth trajectory in low K^+^ media (Epstein et al., 1993). We transformed TK2463 bacteria with our constructs of interest and controls; expression of the proteins was driven by the IPTG (Isopropyl β-D-1-thiogalactopyranoside)-inducible tac-promoter. In low K^+^-containing medium (∼2 mM K^+^), expression of full length *GmSALT3* improved growth compared to expression of truncated *Gmsalt3*, and the negative control (AtCBL1n, a 12 aa peptide with the sequence MGCFHSKAAKEF; Batistič, Waadt, Steinhorst, Held, & Kudla, 2010) (Fig. 1a; 1b). Arabidopsis AtCHX20 (Chanroj et al., 2011) and AtKAT1 (Anderson, Huprikar, Kochian, Lucas, & Gaber, 1992), a known plasma membrane potassium channel from Arabidopsis, were included as positive controls and restored bacteria growth as expected (Fig. 1a; 1b). We had previously used GmSALT3-YFP for determining the subcellular localisation of GmSALT3 (Guan et al., 2014), and therefore investigated if this translational fusion is functional and could complement TK2463 *E. coli*. Additionally, we were interested whether the last transmembrane domain of GmSALT3 is important for functionality - truncated *Gmsalt3* from NIL-S contains only 9 out of the 10 predicted alpha-helices. We therefore constructed GmSALT3-TM10, that has all 10 TM alpha -helices with only the large (383 aa) hydrophilic C-terminus removed. Both, YFP-tagged GmSALT3 and GmSALT3-TM10 increased *E. coli* growth to that of GmSALT3, and significantly greater than that of Gmsalt3 (Fig. 1a; 1b). None of the test constructs or controls had a significant impact on bacterial growth rates on high K^+^ media (Fig. S1c-d); or showed significant differences on low K^+^ media without IPTG-induction (Fig. S1a-b). In addition, we measured K^+^ content of *E. coli* cells, to determine the impact of *GmSALT3* expression on K^+^ accumulation. Bacteria were grown for 12 h in low K^+^ following IPTG induction, and then harvested. Results indicated that expression of full length *GmSALT3* leads to higher K^+^ content in bacteria, as does expression of *GmSALT3-YFP, GmSALT3_TM10, AtKAT1*, and *AtCHX20* (Fig. S1e). Expression of truncated *Gmsalt3* resulted in similar low K^+^ content as in the negative control.

**Figure 1.**
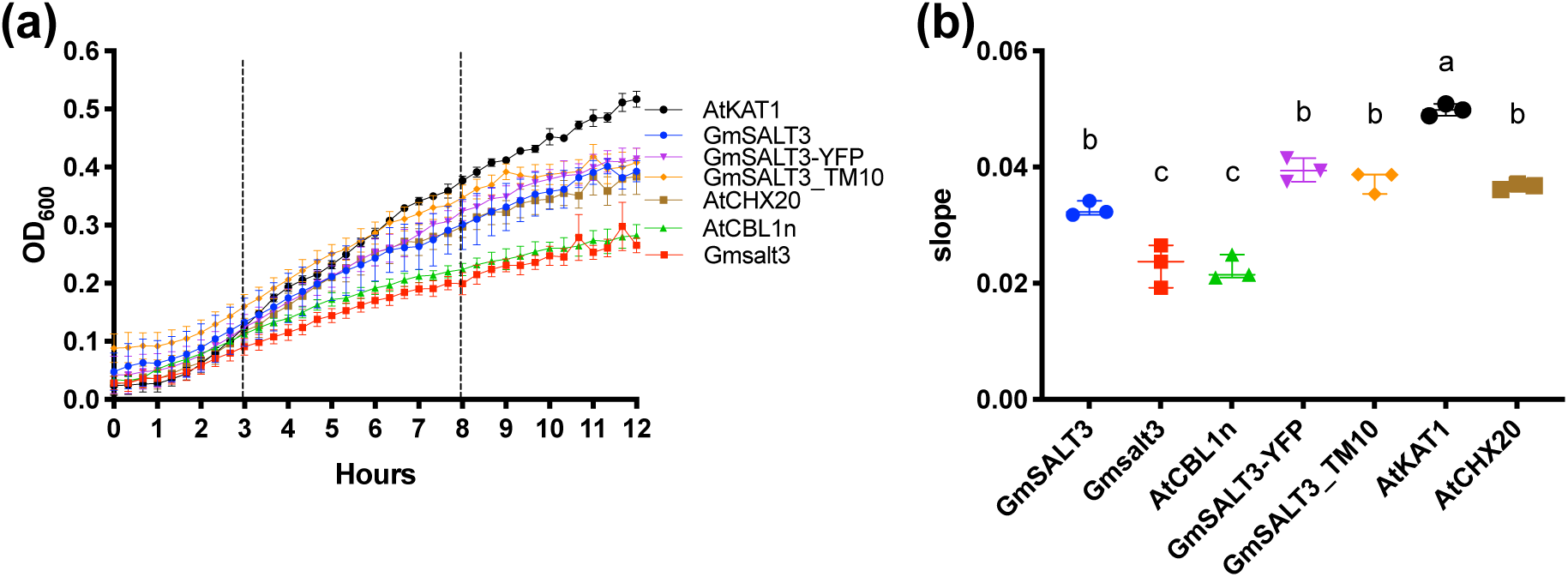
Functional test of *GmSALT3* in *E. coli*. *E. coli* strain TK2463 (defective in potassium uptake, Trk, Kup, and Kdp) harbouring a modified pET-DEST42 vector with *GmSALT3, Gmsalt3, CBL1n, GmSALT3-YFP, GmSALT3-TM10, AtCHX20 and AtKAT1* were grown in different media. (**a)** Low potassium medium (10 g/L tryptone, 2 g/L yeast extract, 100 mM D-mannitol; ∼2 mM K^+^) with additional 0.5 mM Isopropyl β- D-1-thiogalactopyranoside (IPTG). All the test media were supplemented with 100 μg/ml carbenicillin. OD_600_ was monitored every 15 min in 96-well microplate reader. n = 3. Growth rates (OD_600_ per hour) of *E. coli* cells within the log phase are indicated between dotted lines, and their slopes are shown in (**b)** (low potassium medium with IPTG). Different letters indicate statistically significant differences between *E. coli* cells harbouring different constructs (one-way ANOVA followed by Fisher’s LSD test, *p* < 0.05).

We chose a second heterologous expression system to investigate if the ER-localised GmSALT3 impacts the fluxes of ions additional to K^+^. We expressed YFP-tagged and untagged GmSALT3, as well as Gmsalt3, in *X. laevis* oocytes and detected fluorescence at the PM in GmSALT3-YFP expressing oocytes, similar to the PM-localised positive control TmHKT1;5-A-YFP (Fig. 2a) (*Triticum monococcum* High-Affinity K^+^ Transporter 1; Munns et al., 2012). This indicates that GmSALT3-YFP localises to the PM in oocyte, which is different from its ER-localisation in plant cells (Guan et al., 2014). It is not uncommon for plant endomembrane proteins to at least partially localise to the PM in frog oocytes (Maurel, Reizer, Schroeder, & Chrispeels, 1993; Henderson et al., 2015); however, this restricts the interpretation of the data regarding the impact of ER-localised transporters on PM ion fluxes.

**Figure 2.**
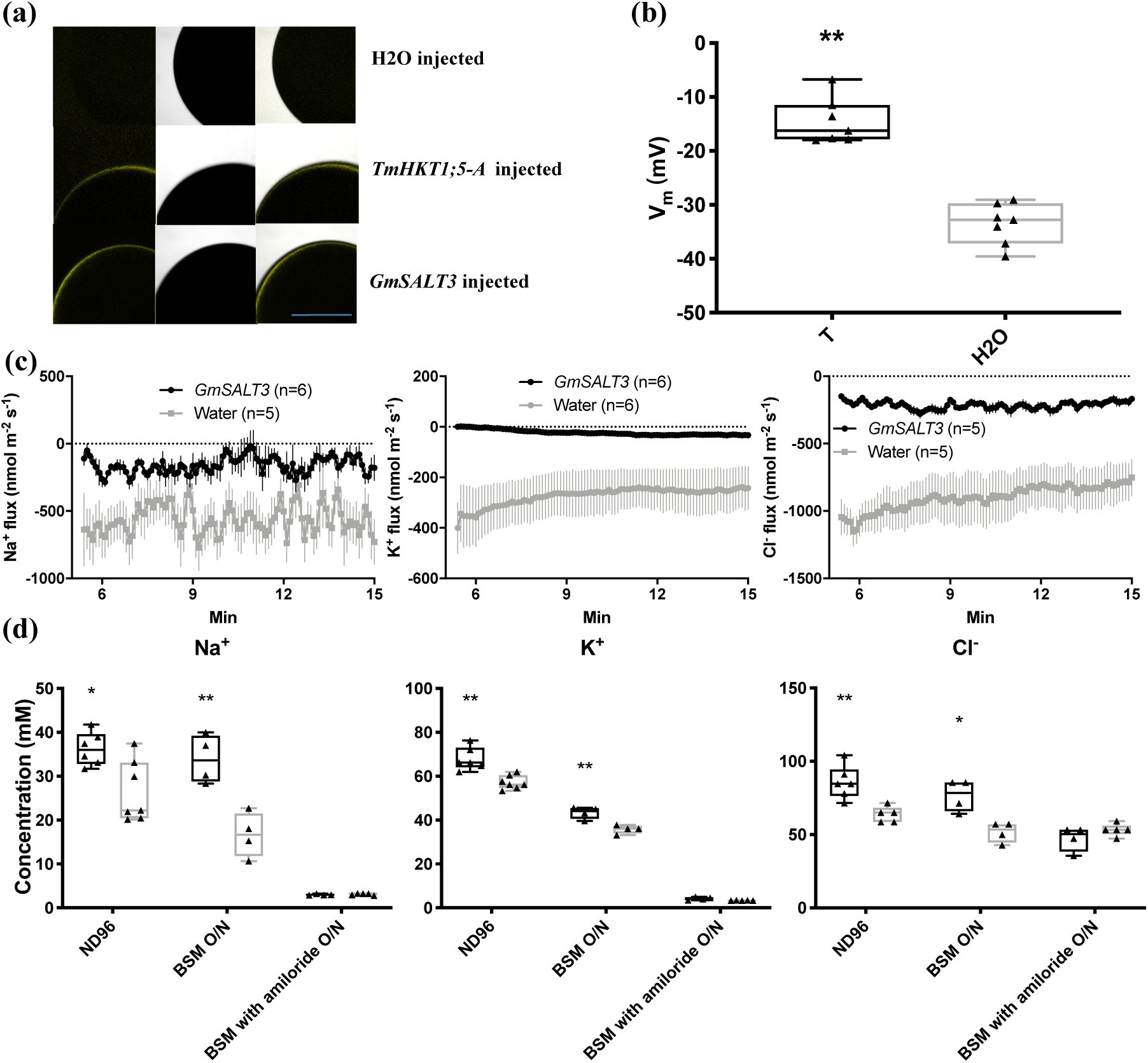
Functional characterization of *GmSALT3* in *X. laevis* oocytes. **(a)** Plasma membrane localisation of expressed *GmSALT3-YFP* in oocytes. H_2_O-injected and *TmHKT1;5-A*-YFP (positive control) expressed oocytes are also shown. Scale bar = 0.5 mm. Oocytes were analysed by confocal laser-scanning microscopy. (**b)** Resting membrane potentials of *GmSALT3*-and H_2_O-injected oocytes after incubation in ND96 for 72 hours. n = 8, Asterisks indicate a significant difference between GmSALT3-and H_2_O-injected oocytes at \**p* < 0.01 according to the LSD test. (**c)** Net ion fluxes were measured on oocytes plasma membrane using MIFE (Microelectrode Ion Flux Estimation). Oocytes were incubated in ND96 for 72 hours, and washed in ND96 in (96 mM NaCl, 1 mM KCl, 1 mM MgCl_2_, 5 mM HEPES, pH 7.5). Measurements were done in BSM (Basal Salt Medium; 5 mM NaCl, 0.2 mM KCl, 0.2 mM CaCl_2_, 5 mM HEPES, pH 7.5) and exchanged after time zero to BSM with sufficiently low concentrations of Na^+^, K^+^ and Cl^−^ to ensure a gradient for efflux for target ions and an improved signal:noise ratio. n = 4 - 6. (**d)** Ion (Na^+^, K^+^, and Cl^−^) concentrations in *GmSALT3* (black) and H_2_O (grey) injected oocytes after incubation in ND96 (72 hours), BSM (overnight), BSM with 100 μM amiloride (overnight). n = 4-7. Asterisks indicate a significant difference between *GmSALT3*-injected and H_2_O-injected oocytes at \**p* <0.05, \*\**p* < 0.01 according to the LSD test.

Two electrode voltage-clamp electrophysiology revealed that the resting membrane potential of GmSALT3-injected oocytes was more positive compared to H_2_O-injected oocytes when incubated in ND96 media (Fig. 2b), but no consistent current differences were detected. This suggests that transport through GmSALT3 might be electroneutral, which is consistent with the CPA2 signature sequence for electroneutral ion transport (Masrati et al., 2018). Therefore, we chose to conduct MIFE (Microelectrode Ion Flux Estimation) experiments to measure the net fluxes of specific ions across the plasma membrane of oocytes non-invasively and so does not rely upon net charge movement in a particular direction.

Flux analysis indicated that in *GmSALT3*-injected oocytes net efflux of Na^+^, K^+^, and Cl^−^ was reduced compared to H_2_O-injected oocytes (Fig. 2c). Subsequent ion content analysis of oocytes revealed that *GmSALT3*-injected oocytes contained significantly more K^+^, Na^+^, and Cl^−^ compared to H_2_O-injected oocytes, in agreement with the reduced net ion efflux in oocytes (Fig. 2d). To test whether the GmSALT3 induced differences in ion concentration could be blocked, we used the Na^+^ channel and Na^+^/H^+^ exchanger inhibitor amiloride hydrochloride (Darley et al., 2000). Oocytes were transferred from ND96 to BSM (low Na^+^, K^+^ and Cl^−^) and incubated overnight with or without the inhibitor. *GmSALT3*-injected oocytes incubated without amiloride, showed a smaller relative K^+^, Na^+^, and Cl^−^ decrease in compared to H_2_O-injected oocytes as expected (Fig. 2d). In the presence of amiloride, however, this difference was not observed (Fig. 2d), strongly suggesting that GmSALT3 affects the transport of K^+^, Na^+^, and Cl^−^ across the PM in oocytes. Collectively, the heterologous data has advanced our knowledge by confirming that *GmSALT3* encodes a competent transport protein, certainly capable of transporting K^+^, and affects the flux of Na^+^, K^+^ and Cl^−^ across a plasma membrane. However, as it is unclear whether the net fluxes of Na^+^, K^+^ and Cl^−^ across the plasma membrane of oocytes were all carried directly by GmSALT3, rather than endogenous *X. laevis* transport proteins we focused our efforts on further characterising the impact of GmSALT3 on salt transport *in planta*.

### GmSALT3 modulates Na^+^, K^+^, and Cl^−^ accumulation in soybean

Liu et al. (2016) previously used an extreme salt treatment (200 mM NaCl) screen that typically results in the death of even the more salt tolerant germplasm after a prolonged exposure. Using this method, it was observed that full-length *GmSALT3* improved shoot Na^+^ and Cl^−^ exclusion over 10 days, with higher shoot Cl^−^ accumulation occurring prior to shoot Na^+^ accumulation in NIL-S (Liu et al., 2016). Here, we used a lower salt concentration (100 mM) that soybean plants are more likely to encounter in the field, and which typically does not result in death of the plants prior to seed production. Furthermore, we studied the time course of Na^+^, Cl^−^ and K^+^ accumulation in NIL-T and NIL-S for different leaf types, stems and hypocotyls (Fig. 3). We confirmed that Cl^−^ accumulates to higher concentrations in NIL-S shoots compared to those of NIL-T (Fig. 3b). Even without a salt treatment Cl^−^ concentration was found to be greater in NIL-S aerial tissues (leaves and stems), whereas significantly higher Na^+^ accumulation was detected in aerial tissues only after 3 days of NaCl treatment in NIL-S, with Cl^−^ being even more pronounced as time after salt treatment progressed (Fig. 3a,b; Fig. S2, S3). We further observed that the K^+^/Na^+^ ratio was significantly higher in NIL-T leaves compared to NIL-S when measured after day 3 (Fig. 3d). Total K^+^ content was increased in NIL-S compared to NIL-T (Fig. 3c; Fig. S3), but K^+^ accumulation increased to lesser extent compared to Na^+^ accumulation. K^+^ increased from 35.7±1.2 mg/g (day 0) to 51.2±1.2 mg/g (day 10), compared to 0.3±0.1 mg/g and 45.3±5.6 mg/g for Na^+^ in the same period (Fig. 3a; 3c). Therefore, although shoot K^+^ increased in NIL-S, the K^+^/Na^+^ ratio was lower. In addition, salt treatment affected biomass production and after 10 days NaCl treatment the total dry weight of NIL-T was significantly higher, as were individual dry weights of leaves, roots, and stem (Fig. 3e, f).

**Figure 3.**
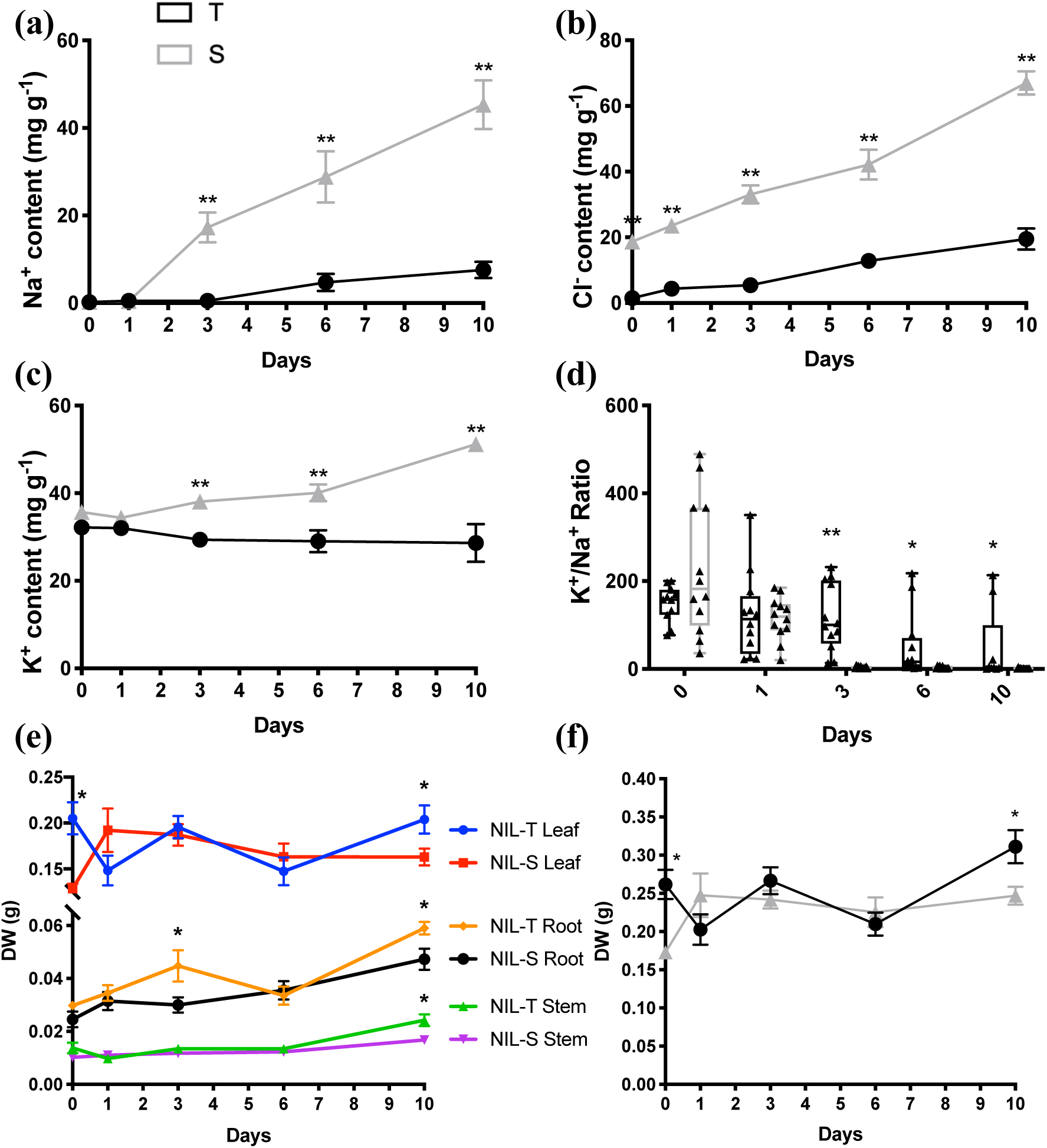
Time-course ion concentration (mg DW^-1^) in NIL-T and NIL-S leaves during 10 days 100 mmol L^-1^ NaCl stress. Na^+^ content (**a**), Cl^**–**^ content (**b**), K^**+**^ (**c**) content, and K^+^ to Na^+^ ratio (**d**) in leaves of NIL-T (black) and NIL-S (grey). Dry weight of different tissues (**e**; comparisons are made between different tissues of NIL-T and NIL-S at different time points) and all tissues (**f**). n = 4. Asterisks indicates a significant difference between NIL-T and NIL-S at \**p* <0.05, \*\**p* < 0.01 according to the LSD test. DW, dry weight.

After establishing the timeline for shoot ion accumulation in 100 mM NaCl, we selected a 4-day time point for further experiments, as the differences in both Na^+^ and Cl^−^ accumulation in NIL-T and NIL-S tissues can be clearly discerned (Fig. 4, Fig. S5), and performed a more detailed analysis of ion concentrations in a range of tissue types, and in the vascular sap. After 4 days, Na^+^ accumulated to a greater extent in all aerial tissues in NIL-S, including leaves (both first trifoliate leaf, FL, and youngest trifoliate leaf, YL), petioles of the first trifoliate leaf (FLP) and youngest trifoliate leaf (YLP), higher stem (HS), lower stem (LS), and hypocotyl (Hy). This is consistent with what we observed for whole shoot ion content analysis. No significant differences were observed between the NILs for Na^+^ in roots under salt treatment (primary root, PR and secondary root, SR; Fig. 4a). Under control conditions (4 days), Na^+^ content very low, but significantly higher in secondary root (SR) of NIL-T compared to NIL-S, with no significant differences in other tissues (Fig. S5). No significant differences between NIL-T and NIL-S were observed for K^+^ under control conditions (Fig. S5). And except for the higher K^+^ accumulation in leaves, no differences were found in any other of the investigated aerial tissues or roots (Fig. 4b).

**Figure 4.**
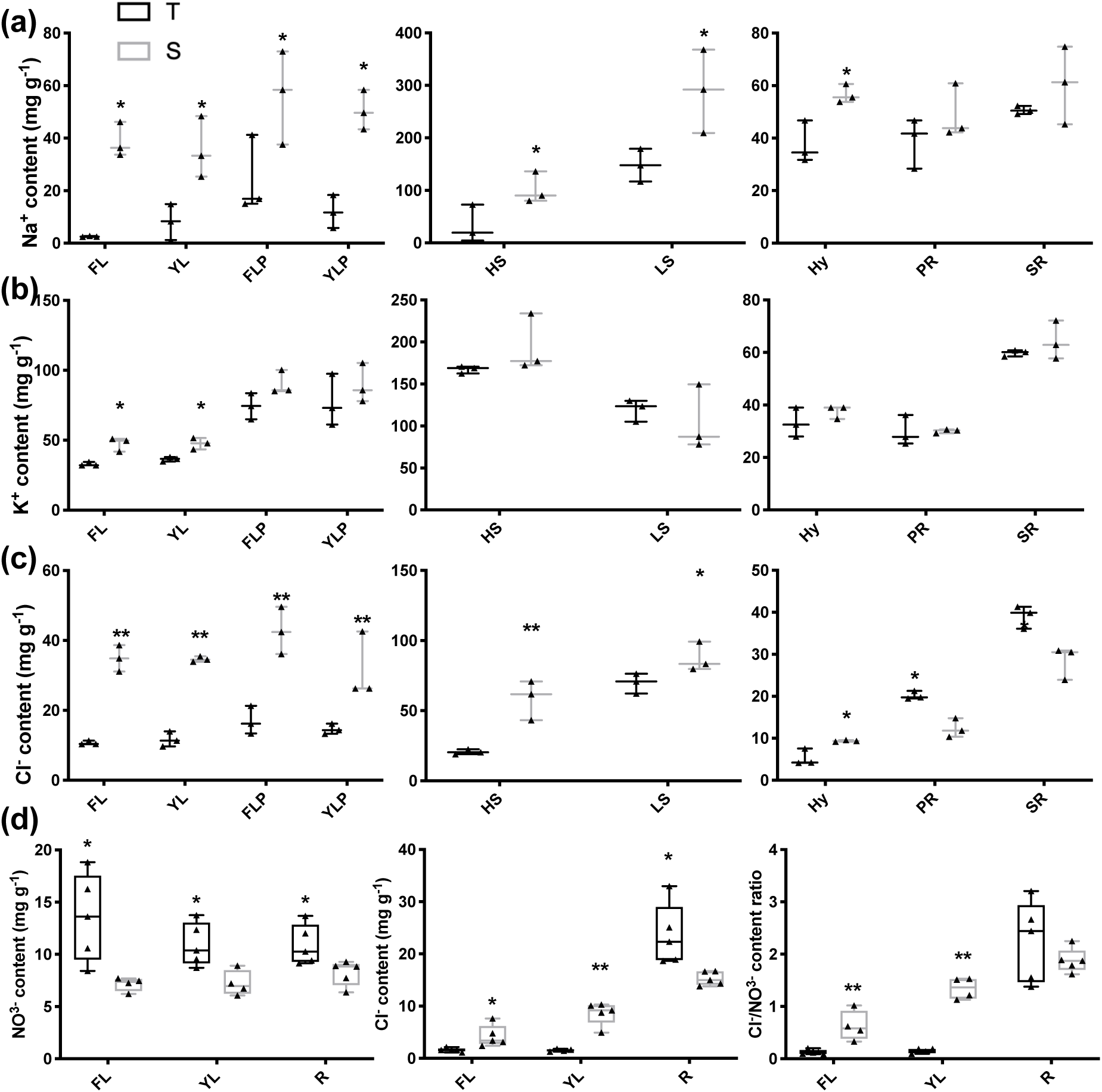
Ion concentration in different tissues of NIL-T and -S after 4 days of 100 mmol L^-1^ NaCl treatment. (**a)** Tissue content of Na^+^ in leaf, hypocotyl and root of NILs. (**b)** K^+^ content in leaf, hypocotyl and root of NILs. (**c)** Cl^−^ content in leaf, hypocotyl and root of NILs. (**d)** NO^3–^ content, Cl^−^ content and Cl^−^/NO^3–^ ratio in leaf and root of NILs. Ion contents are expressed as tissue dry weight. n = 3-5. Asterisks indicate a significant difference between NIL-T (black) and NIL-S (grey) at \**p* <0.05, \*\**p* < 0.01 according to the LSD test (ion content data in control plants with water supply can be found in Figure S4). FL, first trifoliate leaves; YL, youngest trifoliate leaves; FLP, first trifoliate leaves petiole; YLP, youngest trifoliate leaves petiole; HS, higher stem; LS, lower stem; PR, primary root; SR, secondary root; Hy, hypocotyl. Data shown in (**a)**, (**b)**, and (**c)** were from one of the two independent experiments with similar results. Data shown in (**d)** were from a separate experiment and repeated twice.

Cl^−^ accumulation pattern showed similarities but also differences to that of Na^+^ accumulation, under saline conditions. In aerial parts and hypocotyls more Cl^−^ accumulated in leaves, stems and hypocotyls of NIL-S compared to NIL-T (Fig. 4c), similar to Na^+^. While we found that in roots (PR and SR), NIL-T accumulated significantly more Cl^−^ than NIL-S (Fig. 4c). Under control conditions, Cl^−^ content was significantly elevated in leaves and stem of NIL-S, with no differences for hypocotyl and roots (Fig. S5). We also measured NO_3_^−^ content and found that NIL-S accumulated less NO_3_^−^ in leaves (both FL and YL) and roots in compared to NIL-T, which lead to a high. Cl^−^ to NO_3_^−^ ratio in leaves of NIL-S, but not in roots (Fig. 4d). In summary, significantly less Na^+^, K^+^, and Cl^−^ accumulated in NIL-T shoots than NIL-S under saline conditions. Leaf ion exclusion is in many plant species achieved by retention of ions in the root, primarily by reducing net ion concentration in the shoot-ward direction solute flow. To investigate if this is also the case in soybean, we determined xylem and phloem sap ion concentrations.

Ion concentrations within soybean stem phloem and xylem sap were examined at the 4-day time point following NaCl treatment. Glutamine contents within the phloem sap was used to normalize ion measurements to account for volume differences measurements, as glutamine concentration within the phloem remains constant throughout the day (Corbesier, Havelange, Lejeune, Bernier, & Périlleux, 2001). It was found that our estimates of Na^+^ and K^+^ contents were not significantly different between the phloem sap of NIL-T and -S plants. As expected, the Na^+^ concentration in the xylem sap in NIL-S was significantly greater compared to NIL-T, indicating less Na^+^ is loaded into the xylem, or more is retrieved. Surprisingly, and in apparent contrast to the leaf accumulation data, the Cl^−^ concentration in xylem sap was lower in NIL-S than NIL-T (Fig. 5b). In addition to the higher xylem sap concentrations, Cl^−^ concentration was also significantly higher in the phloem sap in NIL-T (Fig. 5a). Phloem sap was harvested from a position where the flow direction is root-wards, close to the junction of root and shoot. The higher Cl^−^ in NIL-T phloem sap therefore indicated an increased flow of Cl^−^ from the shoot to the roots in NIL-T plants, this would be consistent with Cl^−^ being recirculated from the stem to the roots via the phloem. Under control conditions (4 days with water treatment), no significant differences were observed between NIL-T and NIL-S for the ion concentrations within stem phloem and xylem sap (Fig. S5d), which may be related to the detection levels of the assays.

**Figure 5.**
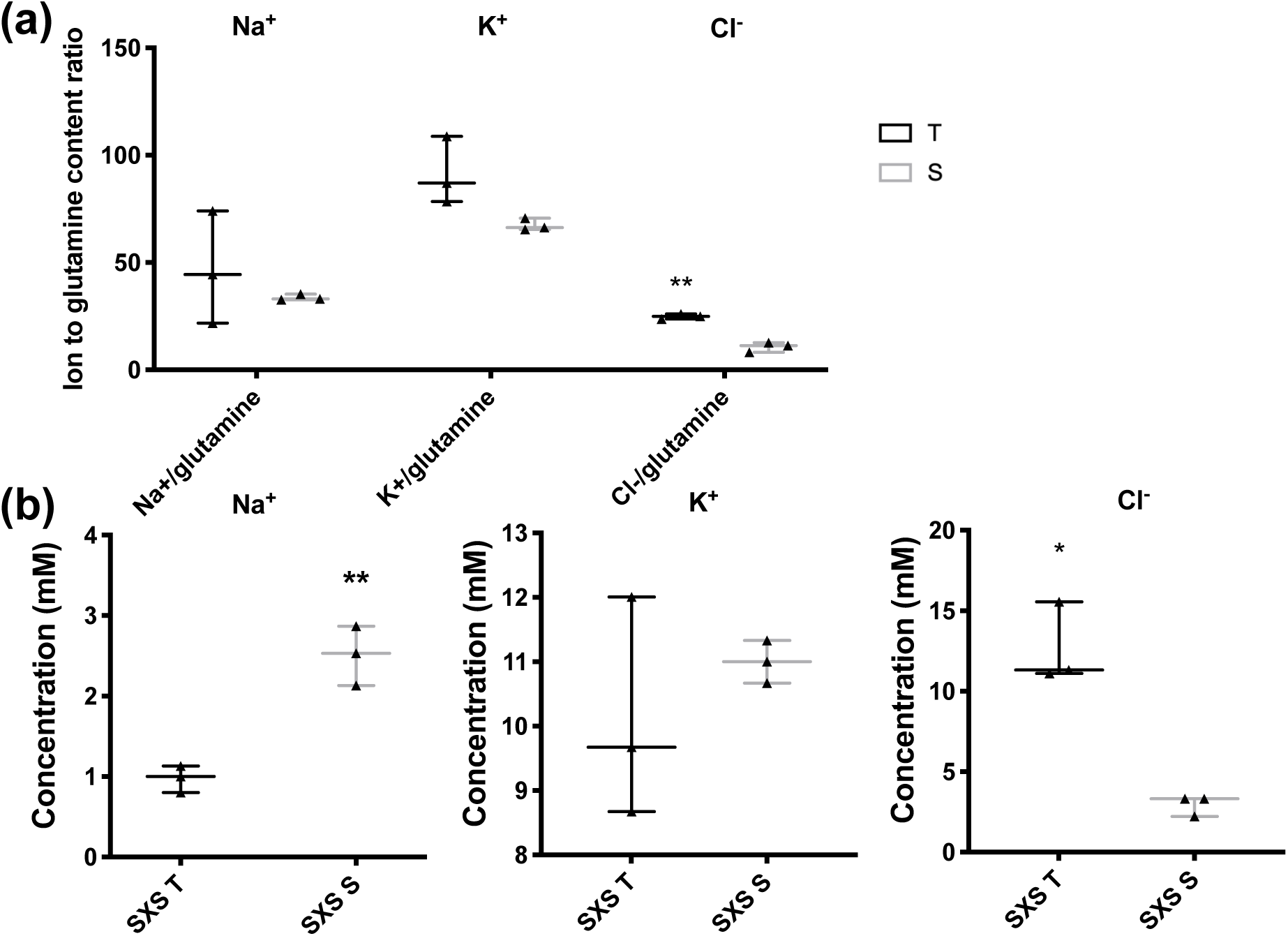
Ion concentration in NIL-T and -S stem phloem and xylem sap after 4 days of 100 mmol L^-1^ NaCl stress. (**a)** Na^+^, K^+^, or Cl^−^ content to glutamine ratio in NIL-T and NIL-S stem phloem sap. (**b)** Na^+^, K^+^, and Cl^−^ concentrations in NIL-T (black) and NIL-S (grey) stem xylem sap. n = 3. Asterisks indicate a significant difference between NIL 820-T and 820-S at \**p* <0.05, \*\**p* < 0.01 according to the LSD test. SPS, stem phloem sap; SXS, stem xylem sap. Data shown here were from one of the three independent experiments with similar results.

The observed higher Cl– concentration in phloem sap suggested that the leaf Cl– exclusion mechanism in NIL-T might be derived from the shoot, instead of the root. To investigate this, a grafting experiment was performed. Reciprocally grafted and self-grafted control plants were salt-treated (100 mM NaCl) for 8 days, as previously described (Guan et al., 2014). The differences in Na^+^ that we observed were consistent with previous grafting experiments, confirming that Na^+^ exclusion depends on a root derived mechanism (Fig. S6a, Guan et al., 2014). In contrast to the situation with Na^+^, grafting demonstrated that the scion, not the rootstock, is important for leaf Cl– exclusion. When a NIL-T scion was grafted on a NIL-S rootstock, the Cl^−^ content in leaves was 89.1% lower when compared to leaves of self-grafted NIL-S (Fig. 6a); with fresh and dry weight calculations showing the same trend (Figure 6b). Conversely, when the NIL-S scion was grafted on the NIL-T rootstock, the Cl^−^ content in leaves was not significantly different compared to self-grafted NIL-S (Fig. 6a). Controls confirmed that self-grafted NIL-S had a considerably higher Cl^−^ content compared to self-grafted NIL-T (Fig. 6a), as expected. In the roots, no significant difference were detected between grafted plants (Fig. 6c).

**Figure 6.**
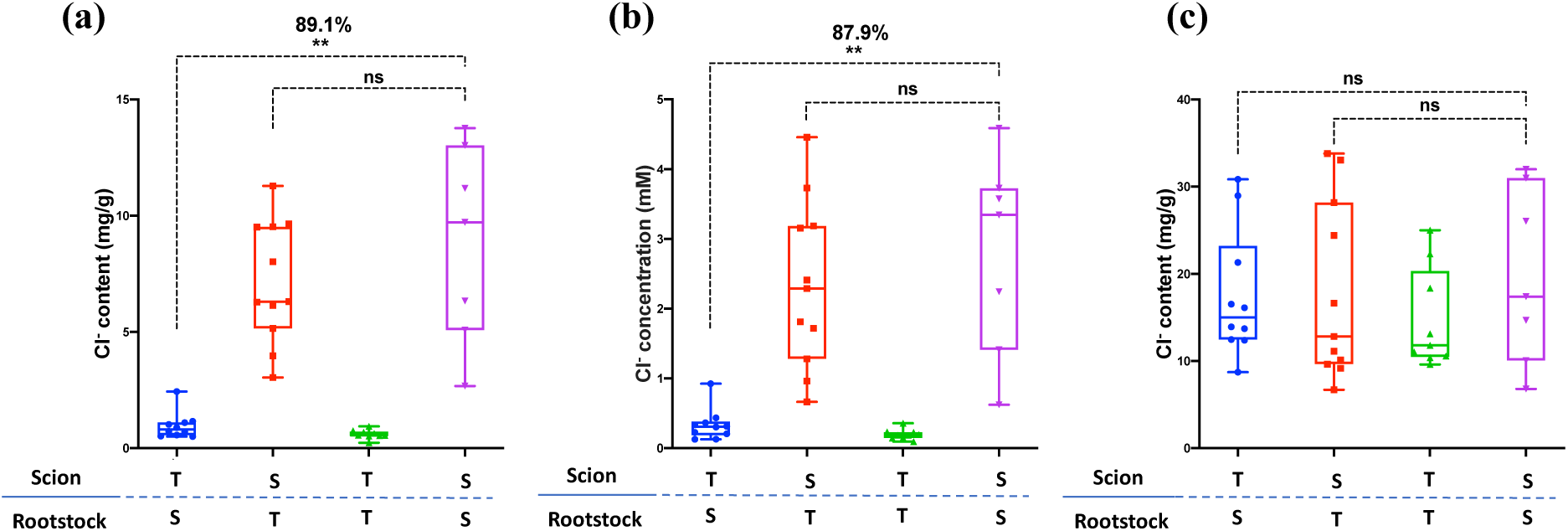
Ion content (Cl^−^) measurement of grafted NIL-T and NIL-S. The leaf Cl^−^ content **(a)** and Cl^−^ concentration **(b)** of reciprocally grafted and self-grafted NIL-T and NIL-S was measured under salt stress (100 mM NaCl) for 8 days. The root Cl^−^ content **(c)** was measured under same conditions. n = 6-11. Asterisks indicate a significant difference between NIL 820-T and 820-S at \**p* <0.05, \*\**p* < 0.01 according to the LSD test. T, NIL-T (*GmSALT3*); S, NIL-S (*Gmsal3*); ns, no significant difference.

As we identified the phloem as an important component of the GmSALT3-derived salt exclusion mechanism, we investigated the ultrastructure of the phloem to identify possible differences between NIL-T and NIL-S. In the phloem, the ER of companion cells reaches into the sieve elements and is directly exposed to the phloem sap (Turgeon & Wolf, 2009). This could theoretically enable a direct ion transport from and into the sap of phloem cells, through the ER-localised GmSALT3, and/or loss of GmSALT3 could lead to morphological changes of the ER ultrastructure. TEM (Transmission Electron Microscopy) imaging did not reveal obvious morphological differences between salt-treated NIL-T and NIL-S in root phloem cells (Fig. S7), indicating lack of full-length *GmSALT3* does not result in a disruption to sub-cellular morphology and that changes in ion flux are more likely directly responsible.

## Discussion

### Transport function of GmSALT3

Plant CPA2 (Cation-Proton Antiporter 2)/CHX (Cation/H^+^ Exchangers) have traditionally been characterised using knockout strains of yeast. A close homolog of GmSALT3, AtCHX20 in *Arabidopsis thaliana*, is preferentially expressed in stomatal guard cells, and shows a localisation pattern consistent with ER-localisation, like GmSALT3 (Padmanaban et al., 2007). Similar to GmSALT3, expression of *AtCHX20* was shown to enhance growth of (a different) *E. coli* strain deficient in K^+^ uptake (LB2003) (Chanroj et al., 2011), indicating that the two proteins catalyse the transport of K^+^. This assay also showed that YFP-tagged GmSALT3 and GmSALT3-TM10 were also functional for K^+^ transport, but not the truncated Gmsalt3, which lacks the last membrane embedded TM10. This suggests that the entire set of transmembrane domains are required for ion transport, and that Gmsalt3 is non-transport competent. Our results further suggest that the large C-terminus, found in all plant CHX proteins, is not required for CHX to conduct ion transport.

We then attempted a transport assay with greater resolution i.e. *X. laevis* oocytes, which has not been previously reported for plant CHX proteins. Using oocytes, two *Drosophila* CPA2 transporters, NHA1 and NHA2, were speculated to act as a Na ^+^/H^+^ exchanger and a H^+^/Cl^−^ cotransporter, respectively (Chintapalli et al., 2015), indicating that proteins from this family can putatively act as either, cation or anion transporters. We observed that GmSALT3 targeted to the PM (Fig. 2a), unlike *in planta*, and led to a lower net ion efflux of all three ions (K^+^, Na^+^ and Cl^−^) across the PM (Fig. 2c). Interestingly, the reduced efflux effect was inhibited by the Na^+^ channel and Na^+^/H^+^ exchanger inhibitor amiloride hydrochloride. Whilst this inhibitor is thought to be selective for Na^+^ transporters, it had an impact on the efflux rate of all three ions measured. It is possible that observed differences in ion flux and accumulation in heterologous systems are the result of changing cytosolic ion concentration that then impacts ion gradients across the PM and influences the activity of native ion transporters. In such a scenario the fluxes detected across the PM of oocytes would be the result of a combination of GmSALT3 and native transporters; and the inhibition of one component leads to the inhibition of the entire mechanism. To conclusively characterise the transport properties of GmSALT3 alternative techniques will need to be employed such as the incorporation of pure protein into liposomes devoid of other transport proteins (Nagarajan et al. 2016).

### The effect of GmSALT3 on ion accumulation *in planta* following salt treatment

Salt tolerance includes tolerance to elevated levels of both Cl^−^ and Na^+^ in the soil solution. In many plant species one of the two ions is understood to be more deleterious to the plant when accumulated in the tissue; however, the situation for soybean is currently unclear with evidence for both ions being more toxic than the other (Läuchli, 1984). Whilst our work does not answer which ion is the most toxic, it does clearly demonstrate that Na^+^ and Cl^−^ long distance transport, and its accumulation in shoots, can be connected to a single gene, which renders the tolerance mechanisms currently inseparable. That said, although leaf Na^+^ and Cl^−^ exclusion are connected to the same gene, *GmSALT3*, confers two distinct mechanisms that are uncoupled *in planta*. Initial evidence suggested that two separate leaf ion exclusion mechanisms exist in soybean comes from our experiments using plants treated with 100 mM NaCl. This salt concentration mimics a more physiologically relevant salt treatment than has previously been used on NIL plants (i.e. one that would be found in the field). Our data suggested that in these conditions shoot exclusion of Cl^−^ broke down prior to exclusion of Na^+^ in NIL-S when compared with NIL-S plants (Fig. 3), and therefore, the mechanisms regulating Cl^−^ and Na^+^ transport are likely to be distinct. More conclusive evidence was highlighted by the reciprocal grafting of NIL-T and NIL-S shoots and roots. These data indicate GmSALT3 is likely to function in the shoot and stem to limit the accumulation of Cl^−^ in the shoot (Fig. 6a). In contrast, Na^+^ was accumulated to greater concentrations in the aerial parts of NIL-S but no difference was observed in the roots compared to NIL-T (Fig. 4a); in stem xylem sap, there was a lower Na^+^ concentration in NIL-T compared to NIL-S but no significant difference in stem phloem sap estimations (Fig. 5). This indicates more Na^+^ is transported in the NIL-S xylem stream. Na^+^ measurements in leaves, xylem sap and the grafting experiment are consistent with a reduced net loading of Na^+^ xylem in NIL-T roots. Root-derived Na^+^ shoot exclusion mechanisms are well known from other plant species, as described in the introduction. Our new finding that shoot Cl^−^ exclusion is a phenotype conferred by the presence of GmSALT3 in the shoots (through grafting) is consistent with our estimations of phloem Cl^−^ content, and we propose is due to phloem recirculation of Cl^−^ back to the roots. Whilst phloem retranslocation of NaCl in saline conditions has been observed before, where in maize approximately 13-36% of the Na^+^ and Cl^-^ imported to leaves through the xylem was retranslocated to the root again in the phloem (Lohaus et al., 2000); no transport protein could be linked to this phenomenon. Furthermore, phloem loading and unloading of Cl^−^ has not been shown in any other plant species to efficiently correlate with salinity tolerance. The role of GmSALT3 in the transfer of Cl^-^ from the xylem into the phloem sap is consistent with its expression in vascular parenchyma cells in the root and shoot (Guan et al., 2014). Further investigation is needed to increase our knowledge on this new tolerance mechanism and the interconnection between xylem and phloem loading for Cl^−^ transport. The prevention of leaf Cl^−^ accumulation, together with the higher NO_3_^−^ content, suggest that NIL-T may maintain increased metabolic and photosynthetic activity in shoots by maintaining a low Cl^−^/NO_3_^−^ ratio, while NIL-S might suffer from Cl^−^ toxicity (Li et al., 2017; Wege, Gilliham & Henderson, 2017).

### Putative *in planta* mechanisms for GmSALT3 regulating long distance ion transport

Coupled to the complexities of identifying the transport properties of GmSALT3 in its native membrane (the endoplasmic reticulum (Guan et al., 2014), and the complex and distinct processes underlying the different salt ion accumulation phenotypes of the two NILs, we propose that GmSALT3 affects processes at the ER and is not directly mediating ion transport across the PM or tonoplast itself – this makes it unlike most other transport proteins identified as important for salt tolerance. Instead, we propose that, *in planta*, the endomembrane localised GmSALT3 influences ion gradients across the PM through currently unknown downstream effects. This might be through impacting the activity of PM-localised transporters, by changing their PM-abundance or changing cytosolic ion supply, or even more complex effects. For example, if the ionic milieu within the ER is affected this could impact the function or sorting of other membrane transporters or regulatory proteins (Bayer et al., 2017), which then impacts PM fluxes; similarly, a difference in the ER-membrane potential could lead to such effects. How GmSALT3 differentially affects the long-distance transport pathways in shoots and roots will now need to be prioritised. Further experiments will need to be carried out to decipher the novel mechanisms imparted by GmSALT3 that are important for salt tolerance. This might enable us to identify similar mechanisms in other (crop) plants.

### Conclusion

To summarize, this work has provided further insights into the salinity tolerance mechanisms imparted by GmSALT3 *in planta* and in heterologous systems. We propose that, in NIL-T, the presence of full-length *GmSALT3* mediates Na^+^ and Cl^−^ exclusion from shoots through restricting net xylem loading of Na^+^, while Cl^−^ is retranslocated from the shoots back into the roots via the phloem. This is the first time that a protein has been shown to confer phloem recirculation of Cl^−^, and that this confers trait improved salt tolerance *in planta*. Our data also demonstrates that GmSALT3 is a transport competent endomembrane localised protein, but fall short of describing the exact cellular mechanisms through which GmSALT3 alters differential transport processes across different cell-types, which is an area that requires further study. RNA-sequencing using NIL-T and NIL-S plants may be beneficial in investigating if GmSALT3 confers salinity tolerance in soybean via influencing transcription, and what distinctive pathways and genes might be significantly changed in NIL-T and NIL-S roots under saline conditions.

## Funding information

Natural Science Foundation of China, grant number 31830066; Scientific Innovation Project of Chinese Academy of Agricultural Sciences; ARC Centre of Excellence funding CE140100008; ARC Fellowships FT130100709, DE160100804 and DE170100346.

### Acknowledgments

This work was funded by the Natural Science Foundation of China, grant number 31830066 (RG); Scientific Innovation Project of Chinese Academy of Agricultural Sciences (LQ). YQ and SH were supported by ARC Centre of Excellence funding awarded to MG (CE140100008). ARC Fellowships supported MG (FT130100709), SW (DE160100804) and JB (DE170100346). We thank Yue (Crystal) Wu for generating the *AtCBL1n* entry vector; Dr. Sunita Ramesh and Assoc. Prof Zhonghua Chen for providing *AtKAT1* construct used in this study; Dr. Gwenda Mayo from Adelaide Microscopy for providing assistance in TEM.

We thank Dr. Yue (Crystal) Wu for generating the *AtCBL1n* entry vector; Sunita Ramesh and Zhonghua Chen for providing *AtKAT1* construct used in this study; CSIRO, Australia for donating *E. coli* strain TK2463. We thank Adelaide Microscopy, especially Gwenda Mayo for the use of TEM and other imaging facilities. This work was funded by the Natural Science Foundation of China, grant number 31830066 (RG). YQ and SH were supported by ARC Centre of Excellence funding awarded to MG (CE140100008). ARC Fellowships supported MG (FT130100709), SW (DE160100804) and JB (DE170100346).

## Author contributions

MG, YQ and SW designed experiments, with input from LQ and RG. LQ and MG directed the project. YQ performed ion accumulation measurement, TEM, IRGA, *E. coli* and oocyte expression, analysed the data and drafted the manuscript. JB performed MIFE in oocytes. SH assisted with experimental design and provided the *E. coli* expression vectors. YQ, MG and SW wrote the manuscript. All authors commented on the manuscript.

## Supporting Information

**Fig. S1. Functional test of *GmSALT3* in *E. coli* with low potassium and KML medium**.

**Fig. S2. Time-course ion concentration (Na^+^) in NIL 820-T and 820-S leaves, stems, hypocotyls, and roots during 10 days NaCl stress**.

**Fig. S3. Time-course ion concentration (K^+^) in NIL 820-T and 820-S leaves, stems, hypocotyls, and roots during 10 days NaCl stress**.

**Fig. S4. Time-course ion concentration (Cl**^−^**) in NIL 820-T and 820-S leaves, stems, hypocotyls, and roots during 10 days NaCl stress**.

**Fig. S5. Ion content and ion flux in NIL 820-T and 820-S different tissues after 4 days in control conditions**.

**Fig. S6 Ion content (Na^+^ and K^+^) measurement of grafted NIL-T and NIL-S**.

**Fig. S7. Ultrastructure of NIL-*GmSALT3* and NIL-*Gmsalt3* root cross sections with 4 days NaCl stress**.

**Table S1. Primers used in this study. Primers were designed and adjusted using Primer3**.

## References

Abel G. (1969). Inheritance of the capacity for chloride inclusion and chloride exclusion by soybeans. Crop Science, 9, 697–698.

Alexandratos N. & Bruinsma J. (2012). World agriculture towards 2030/2050: the 2012 revision: ESA Working paper No.12-03. Rome, FAO.

Anderson J.A., Huprikar S.S., Kochian L.V., Lucas W.J. & Gaber R.F. (1992). Functional expression of a probable Arabidopsis thaliana potassium channel in Saccharomyces cerevisiae. Proceedings of the National Academy of Sciences, 89, 3736–3740.

Batistič O., Waadt R., Steinhorst L., Held K. & Kudla J. (2010). CBL-mediated targeting of CIPKs facilitates the decoding of calcium signals emanating from distinct cellular stores. The Plant Journal, 61, 211–222.

Bayer E.M., Sparkes I., Vanneste S. & Rosado A. (2017). From shaping organelles to signalling platforms: the emerging functions of plant ER–PM contact sites. Current Opinion in Plant Biology 40, 89–96.

Berthomieu P., Conéjéro G., Nublat A., Brackenbury W.J., Lambert C., Savio C., Uozumi N., Oiki S., Yamada K. & Cellier F. (2003). Functional analysis of AtHKT1 in Arabidopsis shows that Na^+^ recirculation by the phloem is crucial for salt tolerance. The EMBO Journal, 22, 2004–2014.

Blanco, F.F., M.V. Folegatti, H.R. Gheyi, and P.D. Fernandes. 2007. Emergence and growth of corn and soybean under saline stress. Scientia Agricola. 64:451–459.

Cao D., Li Y., Liu B., Kong F., & Tran L.S.P. (2018). Adaptive mechanisms of soybean grown on salt-affected soils. Land Degradation & Development, 29, 1054–1064.

Chang R.Z., Chen Y.W., Shao G.H., Wan C.W. (1994). Effect of salt stress on agronomic characters and chemical quality of seeds in soybean. Soybean Science, 13, 101–105.

Chanroj S., Lu Y., Padmanaban S., Nanatani K., Uozumi N., Rao R. & Sze H. (2011). Plant-specific cation/H^+^ exchanger 17 and its homologs are endomembrane K^+^ transporters with roles in protein sorting. Journal of Biological Chemistry, 286, 33931–33941.

Chintapalli V.R., Kato A., Henderson L., Hirata T., Woods D.J., Overend G., Davies S.A., Romero M.F. & Dow J.A. (2015). Transport proteins NHA1 and NHA2 are essential for survival, but have distinct transport modalities. Proceedings of the National Academy of Sciences, 112, 11720–11725.

Corbesier L., Havelange A., Lejeune P., Bernier G. & Périlleux C. (2001). N content of phloem and xylem exudates during the transition to flowering in Sinapis alba and Arabidopsis thaliana. Plant, Cell & Environment, 24, 367–375.

Czerny D.D., Padmanaban S., Anishkin A., Venema K., Riaz Z. & Sze H. (2016). Protein architecture and core residues in unwound α-helices provide insights to the function of a plant transporter AtCHX17. Biochimica et Biophysica Acta (BBA)-Biomembranes, 1858, 1983–1998.

Darley, C.P., Van Wuytswinkel, O.C., Van der woude, K. and Mager, W.H. (2000). Arabidopsis thaliana and Saccharomyces cerevisiae NHX1 genes encode amiloride sensitive electroneutral Na^+^/H^+^ exchangers. Biochemical Journal, 351, 241–249.

Do T.D., Chen H., Hien V.T.T., Hamwieh A., Yamada T., Sato T., Yan Y., Cong H., Shono M. & Suenaga K. (2016). Ncl synchronously regulates Na^+^, K^+^, and Cl^−^ in soybean and greatly increases the grain yield in saline field conditions. Scientific Reports, 6, 19147.

Epstein W., Buurman E., McLaggan D. & Naprstek J. (1993). Multiple mechanisms, roles and controls of K^+^ transport in Escherichia coli: Portland Press Limited.

FAO. (2017). The future of food and agriculture – Trends and challenges, Rome: FAO.

Gilliham M., Able J.A. & Roy S.J. (2017). Translating knowledge about abiotic stress tolerance to breeding programmes. The Plant Journal. 90, 898–917.

Guan R., Qu Y., Guo Y., Yu L., Liu Y., Jiang J., Chen J., Ren Y., Liu G. & Tian L. (2014). Salinity tolerance in soybean is modulated by natural variation in GmSALT3. The Plant Journal, 80, 937–950.

Guo J., Zhou Q., Li X., Yu B. & Luo Q. (2017). Enhancing NO_3^-^_supply confers NaCl tolerance by adjusting Cl^-^ uptake and transport in G. max & G. soja. Journal of Soil Science and Plant Nutrition, 17, 194–202.

Ha B.K., Vuong T.D., Velusamy V., Nguyen H.T., Shannon J.G. & Lee J.D. (2013). Genetic mapping of quantitative trait loci conditioning salt tolerance in wild soybean (Glycine soja) PI 483463. Euphytica. 193, 79–88.

Hamwieh A., Tuyen D.D., Cong H., Benitez E.R., Takahashi R. & Xu, D.H. (2011). Identification and validation of a major QTL for salt tolerance in soybean. Euphytica, 179, 451–459.

Hamwieh A. & Xu D. (2008). Conserved salt tolerance quantitative trait locus QTL) in wild and cultivated soybeans. Breeding Science, 58, 355–359.

Ikeda M. (2005). Distribution of K^+^, Na^+^ and Cl^-^ in root and leaf cells of soybean and cucumber plants grown under salinity conditions. Soil Science and Plant Nutrition, 51, 1053–1057.

Lee G.J., Carter J.T.E., Villagarcia M.R., Li Z., Zhou X., Gibbs M.O. & Boerma H.R. (2004). A major QTL conditioning salt tolerance in S-100 soybean and descendent cultivars. Theoretical and Applied Genetics, 109, 1610–1619.

Lenis J., Ellersieck M., Blevins D., Sleper D., Nguyen H., Dunn D., Lee J. & Shannon J. (2011). Differences in ion accumulation and salt tolerance among Glycine accessions. Journal of Agronomy and Crop Science. 197, 302–310.

Li B., Tester M. & Gilliham M. (2017). Chloride on the Move. Trends in Plant Science, 22, 236–248.

Li W.Y., Wong F.L., Tsai S.N., Phang T.H., Shao G. & Lam H.M. (2006a). Tonoplast-located GmCLC1 and GmNHX1 from soybean enhance NaCl tolerance in transgenic bright yellow (BY)-2 cells. Plant Cell & Environment, 29, 1122–1137.

Li X.J., An P., Inanaga S., Eneji A.E. & Tanabe K. (2006b). Salinity and defoliation effects on soybean growth. Journal of Plant Nutrition, 29, 1499–1508.

Liu Y., Yu L., Qu Y., Chen J., Liu X., Hong H., Liu Z., Chang R., Gilliham M., Qiu L. & Guan R. (2016). GmSALT3, which confers improved soybean salt tolerance in the field, increases leaf Cl^-^ exclusion prior to Na^+^ exclusion but does not improve early vigor under salinity. Frontiers in Plant Science, 7, 1485

Lohaus G., Hussmann M., Pennewiss K., Schneider H., Zhu J.J. & Sattelmacher B. (2000). Solute balance of a maize (Zea mays L.) source leaf as affected by salt treatment with special emphasis on phloem retranslocation and ion leaching. Journal of Experimental Botany, 51, 1721–1732.

Luo G.Z., Wang H.W., Huang J., Tian A.G., Wang Y.J., Zhang J.S. & Chen S.Y. (2005a). A putative plasma membrane cation/proton antiporter from soybean confers salt tolerance in Arabidopsis. Plant Molecular Biology, 59, 809–820.

Luo Q., Yu B. & Liu Y. (2005b). Differential sensitivity to chloride and sodium ions in seedlings of Glycine max and G. soja under NaCl stress. Journal of Plant Physiology, 162, 1003–1012.

Läuchli A. (1984). Salt exclusion: An adaptation of legumes for crops and pastures under saline conditions. In Salinity Tolerance in Plants: Strategies for Crop Improvement (Staples, R.C. and Toenniessen, G.H. eds). New York: Wiley, pp. 171–187.

Masrati G., Dwivedi M., Rimon A., Gluck-Margolin Y., Kessel A., Ashkenazy H., Mayrose I., Padan E. and Ben-Tal N. (2018). Broad phylogenetic analysis of cation/proton antiporters reveals transport determinants. Nature communications, 9, 4205.

Maurel C., Reizer J., Schroeder J.I. & Chrispeels M.J.. (1993). The vacuolar membrane protein gamma-TIP creates water specific channels in Xenopus oocytes. The EMBO Journal, 12, 2241–2247.

Munns R., Day D.A., Fricke W., Watt M., Arsova B., Barkla B.J., Bose J., Byrt C.S., Chen Z.-H., Foster K.J., Gilliham M., Henderson S.W., Jenkins C.L.D., Kronzucker H.J., Miklavcic S.J., Plett D., Roy S.J., Shabala S., Shelden M.C., Soole K.L., Taylor N.L., Tester M., Wege S., Wegner L.H. and Tyerman S.D. (2020), Energy costs of salt tolerance in crop plants. New Phytologist, 225, 1072–1090.

Munns R. & Gilliham M. (2015). Salinity tolerance of crops–what is the cost? New Phytologist, 208, 668–673.

Munns R., James R.A., Xu B., Athman A., Conn S.J., Jordans C., Byrt C.S., Hare R.A., Tyerman S.D. & Tester M. (2012). Wheat grain yield on saline soils is improved by an ancestral Na^+^ transporter gene. Nature Biotechnology, 30, 360–364.

Møller I.S., Gilliham M., Jha D., Mayo G.M., Roy S.J., Coates J.C., Haseloff J. & Tester M. (2009). Shoot Na^+^ exclusion and increased salinity tolerance engineered by cell type-specific alteration of Na^+^ transport in Arabidopsis. Plant Cell, 21, 2163–2178.

Nagarajan Y., Rongala, J., Luang S., Singh A., Shadiac N., Hayes J., Sutton T., Gilliham M., Tyerman S.D., McPhee G., Voelcker N.H., Mertens H.D.T., Kirby N.M., Lee J.-G., Yingling Y.G., Hrmova M., 2016. A barley efflux transporter operates in a Na^+^-dependent manner, as revealed by a multidisciplinary platform. Plant Cell 28, 202–218.

Newman, I. (2001). Ion transport in roots: measurement of fluxes using ion-selective microelectrodes to characterize transporter function. Plant, Cell & Environment 24, 1–14.

Obermeyer G. & Tyerman S.D. (2005). NH_4_^+^currents across the peribacteroid membrane of soybean. Macroscopic and microscopic properties, inhibition by Mg^2 +^, and temperature dependence indicate a SubpicoSiemens channel finely regulated by divalent cations. Plant Physiology, 139, 1015–1029.

Padmanaban S., Chanroj S., Kwak J.M., Li X., Ward J.M. & Sze H. (2007). Participation of endomembrane cation/H^+^ exchanger AtCHX20 in osmoregulation of guard cells. Plant Physiology, 144, 82–93.

Phang T.H., Shao G. & Lam H.M. (2008). Salt tolerance in soybean. Journal of Integrative Plant Biology, 50, 1196–1212.

Phang T.H., Shao G., Liao H., Yan X. & Lam H.M. (2009). High external phosphate (Pi) increases sodium ion uptake and reduces salt tolerance of ‘Pi-tolerant’soybean. Physiologia Plantarum, 135, 412–425.

Qi X., Li M.W., Xie M., Liu X., Ni M., Shao G., Song C., Yim A.K.Y., Tao Y., Wong F.L. & Isobe S. (2014). Identification of a novel salt tolerance gene in wild soybean by whole-genome sequencing. Nature Communications, 5, 4340.

Qiu J., Henderson S.W., Tester M., Roy S.J. & Gilliham M. (2016). SLAH1, a homologue of the slow type anion channel SLAC1, modulates shoot Cl^−^ accumulation and salt tolerance in Arabidopsis thaliana. Journal of Experimental Botany, 67, 4495–4505.

Ren Z.H., Gao J.P., Li L.G., Cai X.L., Huang W., Chao D.Y., Zhu M.Z., Wang Z.Y., Luan S. & Lin H.X. (2005). A rice quantitative trait locus for salt tolerance encodes a sodium transporter. Nature Genetics, 37, 1141–1146.

Roy S.J., Negrão S. & Tester M. (2014). Salt resistant crop plants. Current Opinion in Biotechnology, 26, 115–124.

Rupassara S.I. (2008). Metabolite profiling of leaves and vascular exudates in soybean grown under free-air concentration enrichment. PhD thesis, University of Illinois at Urbana-Champaign.

Rus A., Baxter I., Muthukumar B., Gustin J., Lahner B., Yakubova E. & Salt D.E. (2006). Natural variants of AtHKT1 enhance Na^+^ accumulation in two wild populations of Arabidopsis. PLoS Genetics, 2, 210.

Shabala S., Shabala L., Bose J., Cuin T. & Newman I. (2013). Ion flux measurements using the MIFE technique. Plant Mineral Nutrients: Methods and Protocols, 171–183.

Shi H., Bressan R. (2006). RNA extraction. Methods in Molecular Biology 323: 345.

Singh G. (2010). The soybean: botany, production and uses. Oxfordshire, UK: CABI Publishing.

Sze H. & Chanroj S. (2018). Plant endomembrane dynamics: Studies of K^+^/H^+^ antiporters provide insights on the effects of pH and ion homeostasis. Plant Physiology, 177, 875–95.

Tester M. & Davenport R. (2003). Na^+^ tolerance and Na^+^ transport in higher plants. Annals of Botany, 91, 503–527.

Tester M. & Langridge P. (2010). Breeding technologies to increase crop production in a changing world. Science, 327, 818–822.

Tian H., Baxter I.R., Lahner B., Reinders A., Salt D.E. & Ward J.M. (2010). Arabidopsis NPCC6/NaKR1 is a phloem mobile metal binding protein necessary for phloem function and root meristem maintenance. The Plant Cell, 22, 3963–3979.

Turgeon R. & Wolf S. (2009). Phloem transport: cellular pathways and molecular trafficking. Annual Review of Plant Biology, 60, 207–221.

Uozumi N., Kim E.J., Rubio F., Yamaguchi T., Muto S., Tsuboi A., Bakker E.P., Nakamura T. & Schroeder J.I. (2000). The Arabidopsis HKT1 gene homolog mediates inward Na^+^ currents in Xenopus laevis oocytes and Na^+^ uptake in Saccharomyces cerevisiae. Plant Physiology, 122, 1249–1260.

Wege S., Gilliham M. & Henderson, S.W. (2017). Chloride: not simply a ‘cheap osmoticum’, but a beneficial plant macronutrient. Journal of Experimental Botany, 68, 3057–3069.

Xu G.H., Magen H., Tarchitzky J. & Kafkafi U. (2000). Advances in chloride nutrition of plants. San Diego: Academic Press Inc., pp. 97–150.

